# Crystal structure of mRNA cap (guanine-N7) methyltransferase E12 subunit from monkeypox virus and discovery of its inhibitors

**DOI:** 10.1101/2023.03.12.532263

**Authors:** De-Ping Wang, Rong Zhao, Hao-Feng Wang, Mei-Yue Wang, Wen-Shu Hu, Meng-Meng Lin, Wen Shu, Ji-Min Cao, Wen Cui, Xin Zhou

## Abstract

In July 2022, the World Health Organization announced monkeypox as a public health emergency of international concern (PHEIC), over 85,000 global cases have been reported currently. However, preventive and therapeutic treatments are very limited. The monkeypox virus (MPXV) E12, an mRNA capping enzyme small subunit, is essential for the methyltransferase activity of RNA capping enzymes of MPXV. Here, we solved a 2.16 Å crystal structure of E12. We also docked the D1 subunit c-terminal domain (D1_CTD_) of vaccinia virus (VACV) with E12 to analyze the critical residues of interface between them. These residues are used for drug screening. The top six compounds are Rutin, Quercitrin, Epigallocatechin, Rosuvastatin, 5-hydroxy-L-Tryptophan, and Deferasirox. These findings may provide insights into the development of anti-MPXV drugs.

## 1. Introduction

Monkeypox is a zoonotic disease caused by the monkeypox virus (MPXV). The MPXV belongs to the family Poxviridae and is a double-stranded DNA virus [1]. The first human monkeypox virus (MPXV) infection was reported in the Democratic Republic of the Congo [2]. Africans remained the only region affected by monkeypox in humans until 2003 [1, 3]. However, a series of monkeypox cases were identified in May 2022 in some European countries like the United Kingdom [4] and new outbreak was determined quickly by health authorities [5]. Afterwards, a public health emergency of international concern was declared by the WHO on July 23, 2022 [6]. The first case of Monkeypox in China was found in Chongqing municipality on September 16, 2022 [7]. As of February, 2023, more than 85,000 confirmed cases of monkeypox virus infection in humans have been reported in 110 locations worldwide. Endemic monkeypox is usually self-limiting [7]. However, monkeypox can also lead to fatal outcomes in humans. The overall fatality rates of different clades of the monkeypox virus change from 1 to 10% [1, 8]. As a result, in-depth studies of some important proteins which are potential targets for drugs against MPXV are warranted.

MPXV replicates in the cytoplasm of host cells [9]. The poxvirus encodes RNA capping enzymes which play significant roles in the life cycle of poxvirus. The 5’ cap of viral mRNA is essential for the initiation of RNA translation, mRNA stability enhancement, high translation efficiency, and immune escape [10–12]. The mRNA capping enzyme of MPXV is composed of two subunits: E1 (845 aa) and E12 (287 aa), which correspond to D1 and D12 in VACV, respectively. These two subunits are responsible for m7GpppRNA synthesis. The E1 subunit contains three catalytic modules, RNA 5’ triphosphatase (RTPase)-guanylyltransferase (GTPase) modules in the N-terminal and (guanine-N7)-methyltransferase (MTase) module in the C-terminal [10]. In the *Poxviridae* family, these three catalytic activities are required for the synthesis of cap-0. In the cap-0 structure, the 5’ guanine base at the N7 position was methylated by MTase. The E12 subunit binds and stimulates the E1 MTase domain. Then, the MPXV 2’O-MTase VP39 adds another methyl group to the 2’O position of the ribose of the first transcribed base. The cap-1 structure is formed in this step [13].

E12 subunit is of great significance in maintaining the high methyltransferase activity of E1 subunit [14]. In addition, E12 protein is essential for MPXV replication [15]. It has been demonstrated that there is no homolog of E12 in any known eukaryotic organism (or viruses other than poxviruses). Therefore, E12 are promising targets for antipoxviral drug discovery. In this study, we solved the apo form of MPXV E12 for the first time. Besides, we analyzed the critical amino acid located at the heterodimer interface between MPXV E1 and E12 (on the basis of superimposition). We also performed virtual screening based on the E1-E12 interface sites for drug screening. The majority of MTase inhibitors target the AdoMet-binding site. This is therefore the first attempt at drug screening based on the interface between catalytic subunit and stimulatory subunit of poxvirus MTase. It may provide structural information from our study in the mechanism of the MPXV mRNA cap N7 methyltransferase and the novel anti-poxvirus drugs in the future.

## 2. Materials and methods

### 2.1. Expression and purification of MPXV E12 protein

The MPXV E12 gene (861 bp) was chemically synthesized, codon optimized, and cloned in the vector pET 28a for expression in *Escherichia coli*. Cells were grown in LB medium supplemented with 100 ug/mL kanamycin at 37 °C to an absorbance of 0.8 at 600 nm. E12 proteins were induced with 1 mM isopropyl β-D-thiogalactoside (IPTG) at 16 °C for 16-20 h. Cells were collected by centrifugation at 4000 rpm and resuspended with buffer including 50 mM Tris-Hcl, pH 8.0, 500 mM NaCl, 5 mM EDTA, 5% glycerol, 50 uM PMSF, 10 ug/mL Rnase A, 10 ug/mL Dnase I, and then lysed by sonication. After being ultracentrifuged at 4 °C, 15,000 rpm for 1 h, MPXV E12 proteins in supernatant were purified using Ni Sepharose (GE) affinity chromatography, anion exchange chromatography, and size exclusion chromatography successively.

### 2.2. Crystallization of MPXV E12 protein

The crystals of MPXV E12 were cultured for one week at 16 °C using the sitting drop. The crystallisation conditions were as follows: 0.1 M sodium chloride, 0.1 M bis-tris propane pH 9.0, 25% w/v polyethylene glycol 1,500. The crystals were then soaked in a cryoprotectant solution containing 25 mM Tris-Hcl, pH 8.0, 150 mM NaCl, 5 mM DTT, and 20% glycerol before being frozen in liquid nitrogen.

### 2.3. Data collection, structure solution, and refinement

Diffraction data were collected at the Shanghai Synchrotron Radiation Facility (SSRF) beamline BL02U1. Table 1 shows the data collection and processing details. Molecular replacement was used to solve the structures using the VACV mRNA MTase D12 structure (PDB: 2VDW) as a research model. PHENIX was used to optimize the model, and the model was built manually utilizing Coot. PyMOL (http://www.pymol.org/) was used to perform the figures.

**Table 1.** Data collection and refinement statistics.

| Data | MPXV E12 |
| --- | --- |
| **Data Collection** |  |
| X-ray Source | SSRF beamline BL02U1 |
| Wavelength (Å) | 0.98322 Å |
| Space group | *P12<sub>1</sub>1* |
| Unit cell a,b,c,α,β,γ (Å) | 53.3, 46.4, 62.5, 90.0, 90.4, 90.0 |
| Resolution range (Å) | 53.25-2.16 (2.28-2.16)<sup>a</sup> |
| Unique reflections | 16534 (2398)<sup>a</sup> |
| Completeness (%) | 100 (100)<sup>a</sup> |
| Redundancy | 6.8 (6.6)<sup>a</sup> |
| *I*/*σI* | 7.6 (2.1)<sup>a</sup> |
| *R*<sub>merge</sub> | 0.139 (0.473)<sup>a</sup> |
| **Refinement** |  |
| Resolution range (Å) | 40.39-2.16 (2.24-2.16)<sup>a</sup> |
| No. of reflections (working/test) | 16493/1629 |
| *R*<sub>work</sub>/*R*<sub>free</sub><sup>c</sup> | 0.200/0.254 (0.256/0.320)<sup>a</sup> |
| Number of atoms |  |
| Protein | 2367 |
| Water | 184 |
| B-factors |  |
| Protein | 26.3 |
| Water | 31.7 |
| r.m.s. deviations |  |
| Bond lengths (Å) | 0.002 |
| Bond angles (°) | 0.43 |
| Ramachandran Plot<sup>d</sup> |  |
| (% favored/allowed/outliers) | 96.14/3.86/0.00 |

<sup>a</sup> Values in parentheses are the statistics for the highest resolution shell.
<sup>b</sup> $R_{\text{merge}} = \sum_{\text{hkl}} \sum_j |\mathbf{I}_{\text{hkl}} - \mathbf{I}_{\text{hkl}}(j)| / \sum_{\text{hkl}} \sum_j \mathbf{I}_{\text{hkl}}(j)$ . $\mathbf{I}_{\text{hkl}}(j)$ and $\mathbf{I}_{\text{hkl}}$ represent the *j*th and mean intensity of reflection hkl.
<sup>c</sup> $R_{\text{work}} = \sum_{\text{hkl}} ||\mathbf{F}_{\text{obs}}| - |\mathbf{F}_{\text{calc}}|| / |\mathbf{F}_{\text{obs}}|$ . $\mathbf{F}_{\text{obs}}$ and $\mathbf{F}_{\text{calc}}$ represent the observed and calculated structure factors, respectively. $R_{\text{free}}$ is the R factor calculated with 10 % of unique reflections as the test set.
<sup>d</sup> The values are reported by PHENIX.

### 2.4. Virtual screening

The 15325 compounds used in this virtual screening experiment were mainly from APExBIO and Selleck compound libraries (https://www.apexbio.cn/screening-library.html, https://www.selleck.cn/). The LigPre module in Estro11.9 platform was used for protonation and energy minimization of all compounds, and the force field was selected as OPLS3e. The crystal structure of MPXV E12 was possessed on Maestro11.9 platform, including the removal of water and ions (retention of Ni ions in the active site), protonation, addition of missing atoms, completion of missing groups, protein energy minimization, energy optimization. The force field was selected as OPLS3e. The selection and refinement of virtual screening was completed by the Glide module in Schrödinger Maestro software. The crystal structure of E12 was possessed by protein preparation wizard module for structural refinement and minimization. All compounds were prepared according to the default settings of the LigPre module. For screening in the Glide module, the prepared E12 subunit were imported. Docking sites were determined based on protein-protein interactions and used as the centroid of the 15 Å box. First, the ligand was redocked to confirm the feasibility of the docking method. The dataset was then screened by SP docking. The SP docking template is suitable for mass screening of compounds. Finally, the XP docking template was used to screen out the pro-ligands with higher scores determined by the SP method.

## 3. Results

### 3.1. Overall structure of the MPXV E12

The MPXV E12 crystals belong to the *P12*_1_1 space group and the diffraction data were collected at 2.16 Å resolution. The relative data and refinement statistics are showed in Table 1. The structure of MPXV E12 comprises six ß-sheets (β1, β2, β3, β4, β5, and β6) and nine α-helices (αA, αB, αC, αD, αE, αF, αG, αH, and αI) (Fig. 1A). The topology structure of the stimulatory subunit E12 exhibits a class I methyl-transferase (MT) like core (Fig. 1B). However, E12 subunit lacks the AdoMet-binding domain, making it devoid of MT activity, which also suggests that E12 does not play a direct role in substrate binding.

**Fig. 1.**
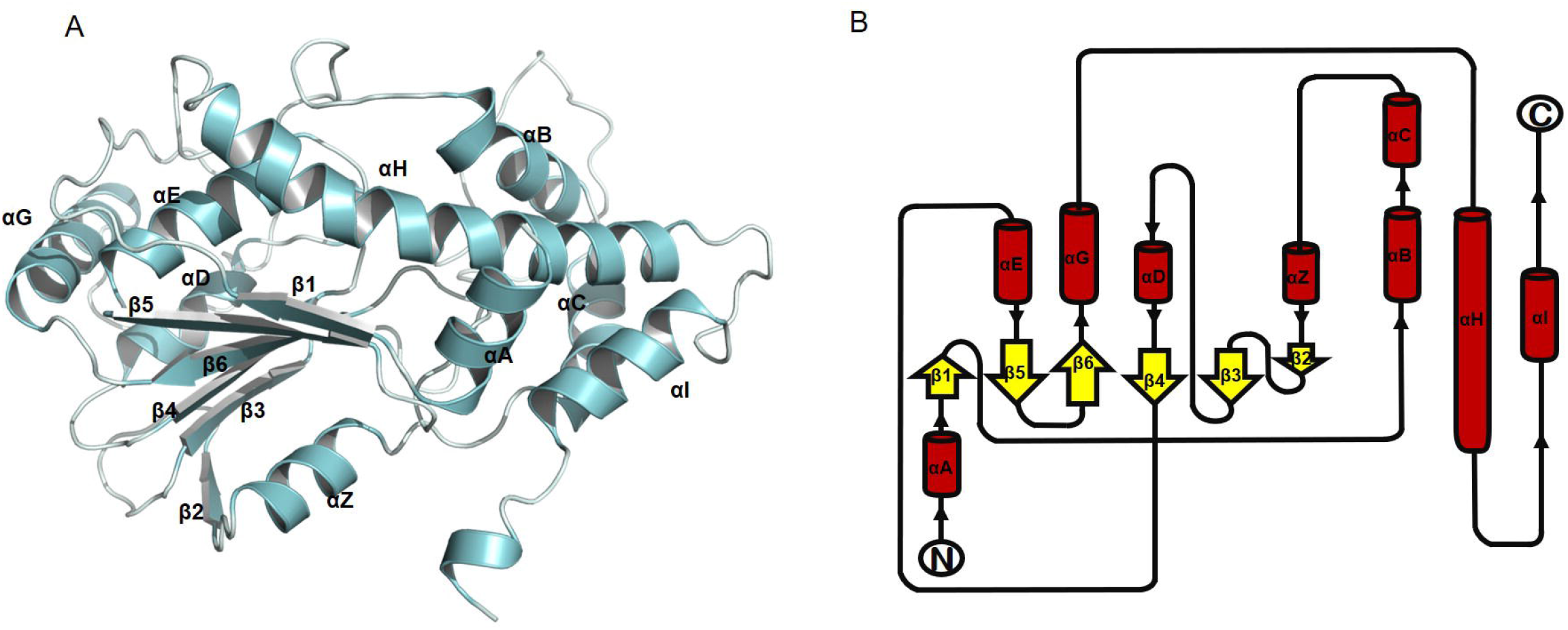
Structure of MPXV E12 subunit. (A) The overall crystal structure of the E12 subunit in cartoon representation. It contains six β-sheets (β1-β6) and nine α-helices (αA, αB, αC, αD, αE, αF, αG, αH, and AI). (B) Schematic diagram of the topology of the E12 subunit.

### 3.2. The E1–E12 interface of MPXV

Instead of having its own MTase activity, E12 may stimulate the weak intrinsic MT activity of E1 on the basis of the study of the VACV stimulatory subunit [16]. We have not obtained the crystal of E1-E12 complex yet. Therefore, we superimposed the crystal structure of D1_CTD_ -D12 complex of the VACV mRNA MTase (2VDW) with E12. The E12 superimposed well with D12. Consequently, The D1_CTD_ and E12 subunits were retained for the generation of heterodimer (Fig. 2A). The Minimization module was used for structural refinement to solve the unreasonable structural contact in protein-protein dock. The force field for energy minimization was selected as OPLS3E, and water molecules were selected for the solvation model. The optimization method was divided into two steps: steepest descent and conjugate gradient, and the maximum number of iterations was 5000.

**Fig. 2.**
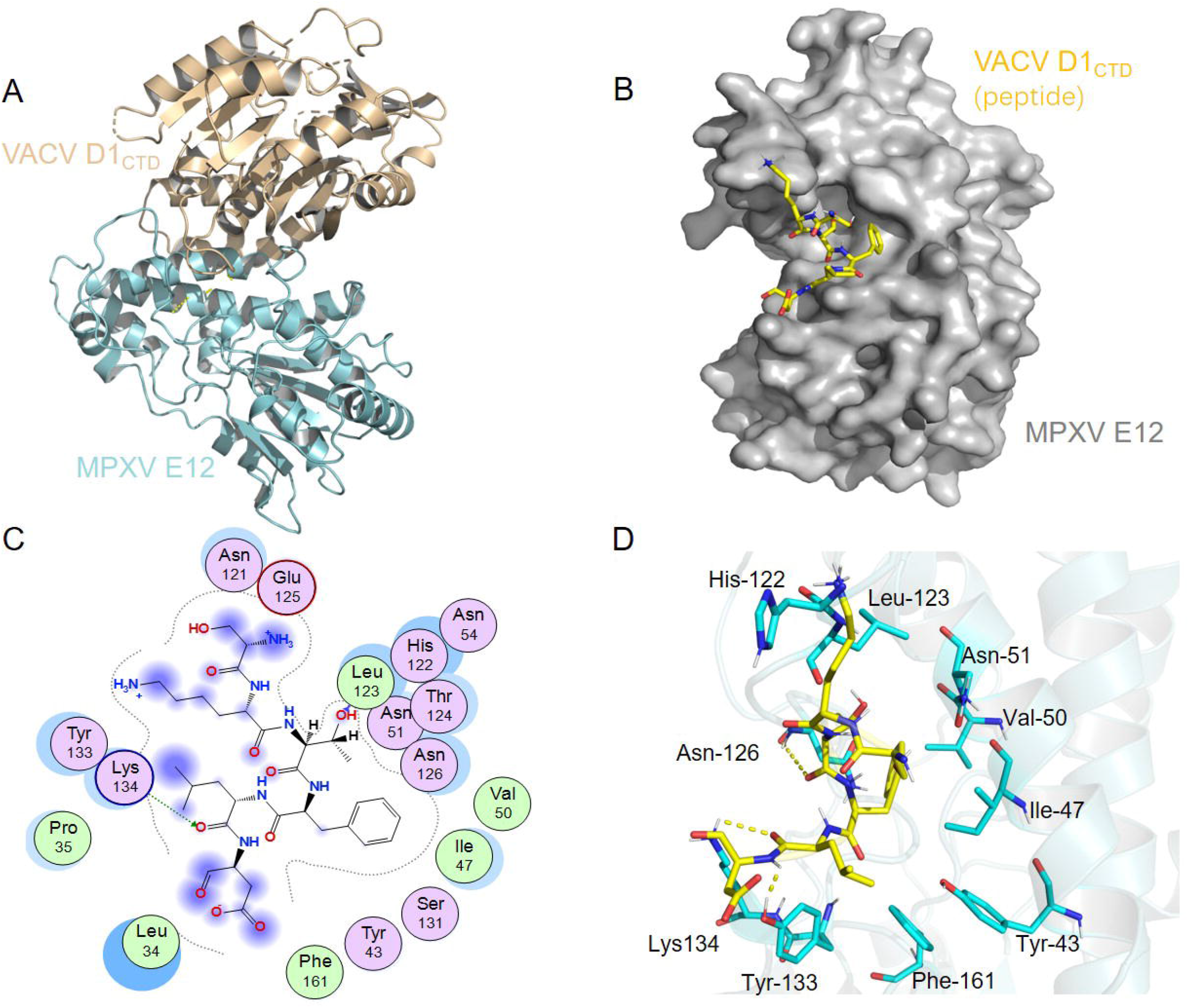
The docking model and analysis of E12 subunit with D1 subunit of vaccinia virus. (A) The overall crystal structure of the D1-E12 complex. The backbone of E12 was rendered in cartoon and colored in cyan. The VACV D1_CTD_ was colored by yellow. (B) A close view of the active site binding with peptide. Key residues interacted with peptide were rendered in stick and colored by yellow. (C) The 2D E12-peptide interaction diagram of complex. E12 residues were rendered in circle and colored based on their properties: green, hydrophobic residue; purple, polar residue. (D) The re-docking results of peptide (yellow) with E12 protein (cyan).

As shown in Fig. S1 and Fig. 2B, a cavity was generated at the binding sites of the two proteins. The amino acids Phe-585 and Leu-586 of D1 inserted into the E12 cavity and strong hydrophobic effect was formed with the cavity. For further confirm the binding sites, D1 protein sequence located in E12 protein cavity was extracted (designated as peptide) as the active site reference and for the conduction of feasibility verification (Fig. 2B). Peptide could form strong hydrogen bond interactions with the key amino acids Tyr-133, Leu-123, Asn-126, and Lys-134 of E12 (Fig. 2C, 2D). Moreover, the hydrophobic residues of the peptide could also form good hydrophobic interactions with Val-50, Ile-47, Tyr-43, Phe-161, and Leu-123 of E12 protein (Fig. 2C, 2D). Previous study shows that Tyr-43 is an absolutely conserved amino acid site and is part of the active domain. These results are in accord with this study [17]. The residues 37-47 in the N-terminal of E12 play an essential role at the interface of E1-E12 and are vital for activity [17].

### 3.3. Inhibitors of the MPXV E12, structure-based virtual screening

The analysis of interface interaction between E12 and D1 based on structure of *MPXV E12* provides a model for identifying inhibitors to target E12 using virtual screening. To achieve this, the ApexBio’s compound library, antiviral compound library, preclinical clinical compound library totaling 15325 compounds, consisting of FDA-approved drugs, clinical-trial drug candidates and natural products, was used for virtual screening of the interface between E12 and D1 sites in the E12 protein (Fig. S2). Based on the docking scores and binding energies, the top 347 compounds were screened out and a list of top 20 potential MPXV E12 inhibitors were shown in Table 2. The top-scoring hits in E1 binding sites of MPXV E12 include Rutin, Quercitrin, Epigallocatechin, Rosuvastatin, 5-hydroxy-L-Tryptophan, and Deferasirox (Fig. S3). These six compounds bind well to the E12 subunit and the matching degrees are high, with binding energy less than −6 kcal/mol. The complexes formed by the docking compounds and E12 are visualized by Pymol 2.1, and the binding modes and critical residues are obtained (Fig. 3A–3F).

**Fig. 3.**
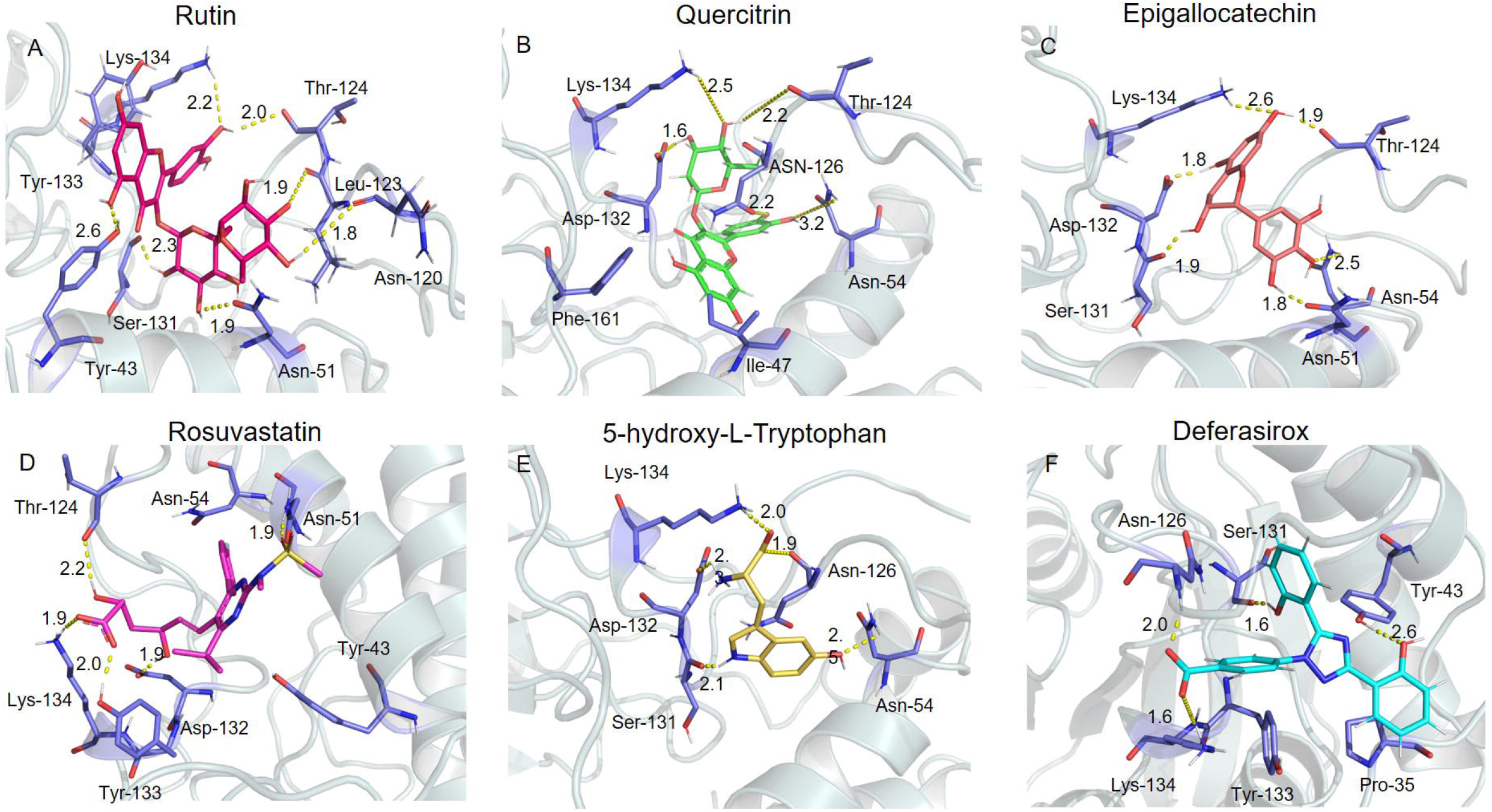
The binding modes of E12 subunit with six compounds. The binding modes of E12 with (A) Rutin, (B) Quercitrin, (C) Epigallocatechin, (D) Rosuvastatin, (E) 5-hydroxy-L-Tryptophan, (F) Deferasirox.

**Table 2.** Virtual screening of compound library for Receptor target.

| Name | CAS | Score | HA | HD | RN | logP | MW |
| --- | --- | --- | --- | --- | --- | --- | --- |
| Rutin | 153-18-4 | -10.35 | 15 | 10 | 6 | -1.113 | 610.521 |
| (-)-Epigallocatechin gallate | 989-51-5 | -8.07 | 10 | 8 | 4 | 3.103 | 458.375 |
| Quercitrin | 522-12-3 | -7.93 | 10 | 7 | 3 | 0.803 | 448.38 |
| Epigallocatechin | 970-74-1 | -7.84 | 7 | 6 | 1 | 1.706 | 306.27 |
| HZ-1157 | 1009734-33-1 | -7.36 | 2 | 0 | 2 | 2.049 | 233.295 |
| Silymarin | 22888-70-6 | -7.11 | 10 | 5 | 4 | 2.641 | 482.441 |
| Rosuvastatin | 287714-41-4 | -7.04 | 6 | 2 | 10 | 1.643 | 480.537 |
| Ceftibuten | 97519-39-6 | -6.57 | 3 | 2 | 7 | -0.37 | 408.415 |
| Pitavastatin Calcium | 147526-32-7 | -6.47 | 3 | 2 | 8 | 4.651 | 420.46 |
| Shikonin | 517-89-5 | -6.42 | 5 | 3 | 3 | 1.277 | 288.299 |
| 5-hydroxy-L-Tryptophan | 4350/9/8 | -6.35 | 1 | 2 | 3 | 1.092 | 220.228 |
| Deferasirox | 201530-41-8 | -6.27 | 4 | 2 | 4 | 3.592 | 372.36 |
| Fluvastatin Sodium | 93957-55-2 | -6.05 | 2 | 2 | 8 | 5.364 | 410.465 |
| Oxytetracycline hydrochloride | 2058-46-0 | -9.16 | 10 | 9 | 2 | -1.457 | 460.439 |
| Hyperoside | 482-36-0 | -8.70 | 11 | 8 | 4 | -0.232 | 464.379 |
| (-)-epigallocatechin | 970-74-1 | -7.84 | 7 | 6 | 1 | 1.706 | 306.27 |
| (-)-Epigallocatechin gallate | 989-51-5 | -7.81 | 10 | 8 | 4 | 3.103 | 458.375 |
| (-)-Epicatechin | 490-46-0 | -7.31 | 6 | 5 | 1 | 1.979 | 290.271 |
| Myricetin (Cannabiscetin) | 529-44-2 | -6.89 | 7 | 6 | 1 | 1.759 | 318.237 |
| Shikonin | 517-89-5 | -6.49 | 5 | 3 | 3 | 1.277 | 288.299 |

Note: Binding energy fuction:

According to the binding modes, the critical residues for compound binding in the protein pocket are clearly identified (Fig. 4A–4F). Rutin interacts with the active sites of E12 protein (Lys-134, Thr-124, Leu-12, Asn-12, Asn-51, Ser1-31, Tyr-43) and hydrogen bond interactions are formed (Fig. 4A). The hydrogen bond distance is short and the binding ability is strong. It is beneficial for Rutin to be anchored in the active sites of E12 to exert its inhibitory effects. The compound Quercitrin can form strong hydrogen bond interactions with the residues Lys-134, Thr-124, Asn-126, Asn-54, and Asp-132, which are conducive to the stability of Quercitrin. Besides, the benzene ring of Quercitrin forms strong hydrophobic interactions with the residues Ile-47 and Phe-161, which also benefit the stabilization of Quercitrin (Fig. 4B). Residues that interact with Epigallocatechin by hydrogen bonds mainly include Lys-134, Thr-124, Asn-54, Asn-51, Ser-131, and Asp-132 (Fig. 4C). Rosuvastatin binds well with E12 and hydrogen bonds are formed between Rosuvastatin and the amino acids Thr-124, Tyr-133, Asp-132, Tyr-43, Asn-51, and Asn-54 of E12. Besides, the carboxyl group of Rosuvastatin can form a salt bridge with residue Lys-134, which is significant for anchoring Rosuvastatin in the E12 protein pocket (Fig. 4D). 5-hydroxy-L-Tryptophan matches well with E12, hydrogen bonds are formed with the residues Asn-126, Asn-54, Ser-131, and Asp-132. In addition, 5-hydroxy-L-Tryptophan forms a salt bridge with Lys-134 to enhance the stabilization of the complex (Fig. 4E). Deferasirox interacts with the residues Tyr-43, Ser-131 by hydrogen bond interactions and the residue Lys-134 by a salt bridge (Fig. 4F). These interactions facilitate the formation of a stable Deferasirox-E12 complex. In conclusion, the above six compounds perform well in docking score and the binding modes with E12 protein, and stable complexes are formed. The binding modes have high similarity. The compounds Rutin, Quercitrin, Epigallocatechin, Rosuvastatin, 5-hydroxy-L-Tryptophan, and Deferasirox are potential active inhibitors against the E12 protein of monkeypox virus.

**Fig. 4.**
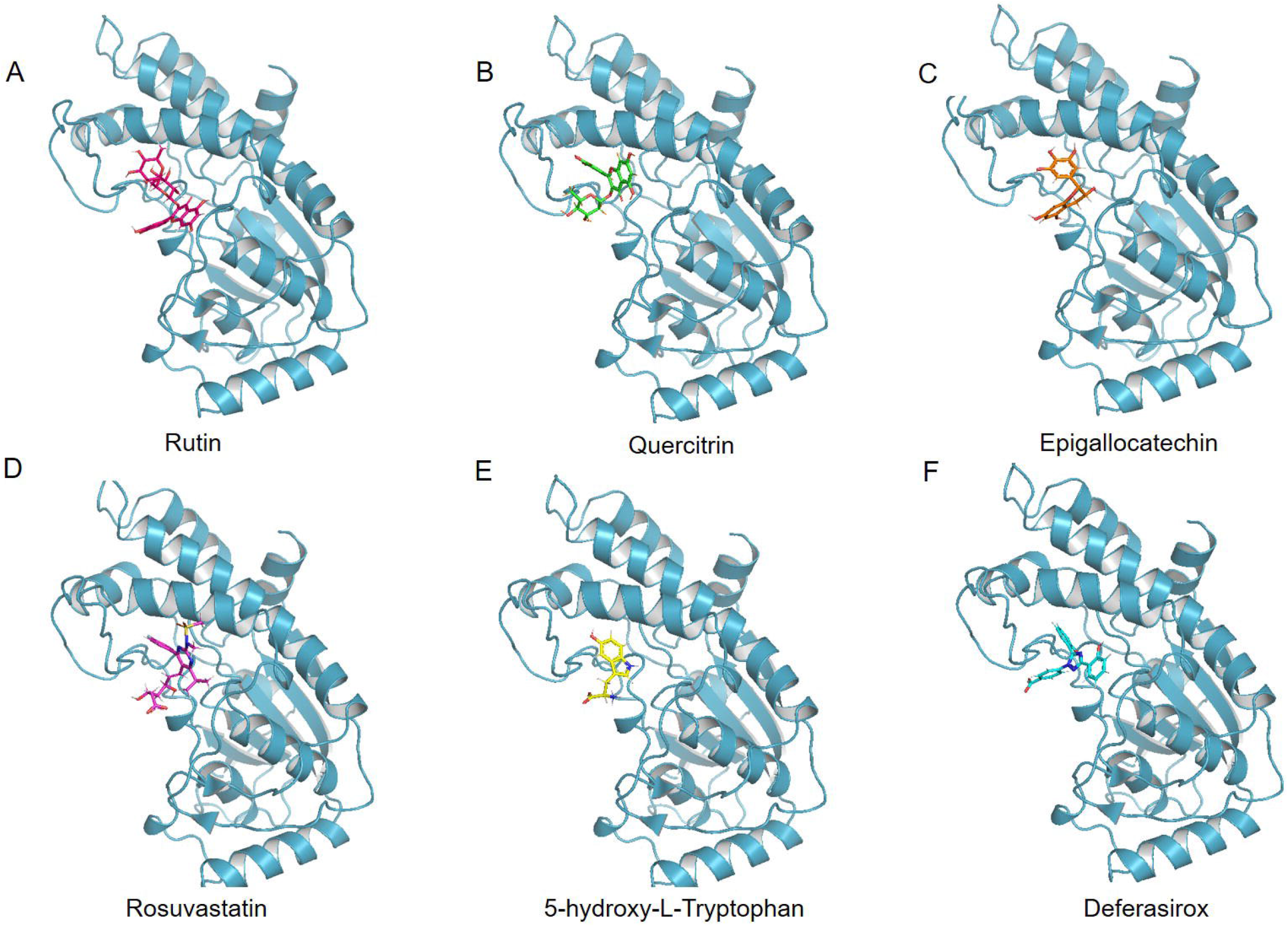
The detailed binding modes of E12 subunit with six compounds. The detail binding modes of E12 with (A) Rutin, (B) Quercitrin, (C) Epigallocatechin, (D) Rosuvastatin, (E) 5-hydroxy-L-Tryptophan, and (F) Deferasirox. The small molecules and the interaction residues were rendered in sticks. Yellow dash represents the hydrogen bond distance or salt bridge.

## 4. Discussion

A global monkeypox outbreak began in 2022, nevertheless, the study about MPXV are limited. Methyltransferase of MPXV plays an important role in expediting efficient viral protein production and escaping from innate immunity detection like other viruses.

The E12 subunit was originally a 2’O-MTase for RNA capping but has gradually evolved to a stimulatory domain for methyltransferase activity enhancement and stabilization of the E1 subunit, allosterically [17]. In previous studies, it was found that the purified carboxyl D1 protein of VACV shows low methyltransferase activity, however, with the addition of purified D12 protein, the methyltransferase activity of the D1 subunit can increase 100-fold [14]. D12 plays an essential role in the replication of VACV but has no substrate binding and catalysis sites for S-adenosyl-homocysteine (AdoHcy) [15]. This may show that although E12 lacks the AdoMet-binding domain and has no methyltransferase activity, it is essential for virus replication and stimulates the MT activity of E1. There is no significant sequence homology between E12 and any other known protein aside from the homologues in the poxviridae family. The E12 subunit is absent even in viruses distantly related to poxvirus-encoding homologs of the E1 subunit. The discovery of antipoxviral drugs based on E12 is therefore promising.

The crystal structure of VACV mRNA capping enzyme shows that the substrate-binding pockets of D1 subunit lie in the opposite direction to the extensive D1-D12 interface [17]. Previous study described the interface between the D12 subunit with VACV MTase and elucidated the possible mechanism of D12 subunit stimulated the MTase activity allosterically. By hydrophobic interactions, the αB’ of D12 subunit packs against the C-terminal part of the MTase helix αK [17]. In addition, the D1-D12 heterodimer also regulate the transcription initiation and termination of intermediate genes and early genes of virus, respectively [18]. These suggest that the amino acid sites at the interface between E12 and E1 play an essential role in methyltransferase function.

In our study, we found Tyr-43, Ile-47, Val-50, Phe-161, Leu-123, Asn-126, Tyr-133, and Lys-134 play significant roles in the virtual screening of E12 inhibitors. The 15,325 compounds were screened and finally 20 compounds were selected by scoring the binding energy and evaluating the critical amino acids of the active site. Top six compounds are Rutin, Quercitrin, Epigallocatechin, Rosuvastatin, 5-hydroxy-L-Tryptophan, and Deferasirox. Rutin is a naturally available flavonoid compound with a wide range of therapeutic effects for many disease [19], such as diabetes [20], cancer [21], inflammatory bowel disease [22], osteoarthritis [23]. (-)-Epigallocatechin gallate (EGCG) is a polyphenol in green tea, with potential therapeutic effects on Parkinson’s disease [24], hypoxia-induced neuroinflammation [25], cancer [26], and so on. Quercitrin is also a natural flavonoid with extensive bioactivities, including antiinflammation, immunomodulation, antioxidative stress, and so on [27]. Rosuvastatin is an HMG-CoA reductase with high efficacy in inhibit hepatic cholesterol synthesis [28]. It is one of the most commonly used stains for the treatment of dyslipidemia and hypercholesterolemia [29, 30]. Besides cholesterol-lowering efficacy, Rosuvastatin also possesses anti-inflammatory, antioxidant and antithrombotic effects [31]. 5-hydroxy-L-Tryptophan (5-HTP) is the immediate precursor in the biosynthetic process of 5-hydroxy-tryptamine (5-HT) [32]. In goat mammary epithelial cells (GMECs), it could elevate the calcium level [33]. Deferasirox is a kind of oral chelator for iron overload. For the treatment of some hematologic disorders, chronic blood transfusions are used, which have the potential to generate iron overload complications. Deferasirox can mobilizes the storage of iron ions in the body by binding with the trivalent form of iron [34–36].

In conclusion, we solved the crystal structure of mRNA cap (guanine-N7) methyltransferase E12 subunit from MPXV and screened its inhibitors by molecular docking. However, the docking results need to be further confirmed by the structure of E1-E12 complex. Besides, cell experiments and virus experiments need to be conducted to confirm the efficacy of the screened compounds.

## Authors contributions

DPW, WC, and JMC designed the study. DPW, RZ, and SW conducted the experiments. HFW, MML and XZ analyzed the data. DPW and MYW drafted the manuscript. WSH and HFW performed the figures. JMC and XZ guided and reviewed the manuscript. All authors read and approved the final manuscript.

## Acknowledgements

This work was supported by Key Medical Science and Technology Program of Shanxi Province (2020XM01), Shanxi “1331” Project Quality and Efficiency Improvement Plan (1331KFC), National Natural Science Foundation of China (81801858, 82170523), and the Applied Basic Research Program of Shanxi Province (202103021224234).

## References

1. Mitjà O., Ogoina D., Titanji B. K., et al., Monkeypox, Lancet. 10370 (401) (2023) 60–74.

2. Di Giulio D. B., Eckburg P. B., Human monkeypox: an emerging zoonosis, Lancet Infect Dis. 1 (4) (2004) 15–25.

3. Beer E. M., Rao V. B., A systematic review of the epidemiology of human monkeypox outbreaks and implications for outbreak strategy, PLoS Negl Trop Dis. 10 (13) (2019) e0007791.

4. Vivancos R., Anderson C., Blomquist P., et al., Community transmission of monkeypox in the United Kingdom, April to May 2022, Euro Surveill. 22 (27) (2022) 2200422.

5. Kluge H., Ammon A., Monkeypox in Europe and beyond-tackling a neglected disease together, Euro Surveill. 24 (27) (2022) 2200482.

6. WHO Director-General’s statement at the press conference following IHR Emergency Committee regarding the multi-country outbreak of monkeypox-23 July 2022 [Available from: https://www.who.int/director-general/speeches/detail/who-director-general-s-statementon-the-press-conference-following-IHR-emergency-committee-regarding-the-multi-country-outbreak-of-monkeypox--23-july-2022.

7. Zhao H., Wang W., Zhao L., et al., The First Imported Case of Monkeypox in the Mainland of China-Chongqing Municipality, China, September 16, 2022, China CDC Wkly. 38 (4) (2022) 853–54.

8. Bunge E. M., Hoet B., Chen L., et al., The changing epidemiology of human monkeypox-A potential threat? A systematic review, PLoS Negl Trop Dis. 2 (16) (2022) e0010141.

9. Peng Q., Xie Y., Kuai L., et al., Structure of monkeypox virus DNA polymerase holoenzyme, Science. 6627 (379) (2023) 100–05.

10. Kyrieleis O. J., Chang J., de la Peña M., et al., Crystal structure of vaccinia virus mRNA capping enzyme provides insights into the mechanism and evolution of the capping apparatus, Structure. 3 (22) (2014) 452–65.

11. Paterson B. M., Rosenberg M., Efficient translation of prokaryotic mRNAs in a eukaryotic cell-free system requires addition of a cap structure, Nature. 5715 (279) (1979) 692–6.

12. Mears H. V., Sweeney T. R., Better together: the role of IFIT protein-protein interactions in the antiviral response, J Gen Virol. 11 (99) (2018) 1463–77.

13. Hodel A. E., Gershon P. D., Shi X., et al., The 1.85 A structure of vaccinia protein VP39: a bifunctional enzyme that participates in the modification of both mRNA ends, Cell. 2 (85) (1996) 247–56.

14. Mao X., Shuman S., Intrinsic RNA (guanine-7) methyltransferase activity of the vaccinia virus capping enzyme D1 subunit is stimulated by the D12 subunit. Identification of amino acid residues in the D1 protein required for subunit association and methyl group transfer, J Biol Chem. 39 (269) (1994) 24472–9.

15. Carpenter M. S., DeLange A. M., A temperature-sensitive lesion in the small subunit of the vaccinia virus-encoded mRNA capping enzyme causes a defect in viral telomere resolution, J Virol. 8 (65) (1991) 4042–50.

16. Saha N., Shuman S., Schwer B., Yeast-based genetic system for functional analysis of poxvirus mRNA cap methyltransferase, J Virol. 13 (77) (2003) 7300–7.

17. De la Peña M., Kyrieleis O. J., Cusack S., Structural insights into the mechanism and evolution of the vaccinia virus mRNA cap N7 methyl-transferase, Embo j. 23 (26) (2007) 4913–25.

18. Vos J. C., Sasker M., Stunnenberg H. G., Vaccinia virus capping enzyme is a transcription initiation factor, Embo j. 9 (10) (1991) 2553–8.

19. Negahdari R., Bohlouli S., Sharifi S., et al., Therapeutic benefits of rutin and its nanoformulations, Phytother Res. 4 (35) (2021) 1719–38.

20. Ghorbani A., Mechanisms of antidiabetic effects of flavonoid rutin, Biomed Pharmacother. (96) (2017) 305–12.

21. Ghanbari-Movahed M., Mondal A., Farzaei M. H., et al., Quercetin-and rutin-based nano-formulations for cancer treatment: A systematic review of improved efficacy and molecular mechanisms, Phytomedicine. (97) (2022) 153909.

22. Habtemariam S., Belai A., Natural Therapies of the Inflammatory Bowel Disease: The Case of Rutin and its Aglycone, Quercetin, Mini Rev Med Chem. 3 (18) (2018) 234–43.

23. Sui C., Wu Y., Zhang R., et al., Rutin Inhibits the Progression of Osteoarthritis Through CBS-Mediated RhoA/ROCK Signaling, DNA Cell Biol. 6 (41) (2022) 617–30.

24. Sergi C. M., Epigallocatechin gallate for Parkinson’s disease, Clin Exp Pharmacol Physiol. 10 (49) (2022) 1029–41.

25. Kim S. R., Seong K. J., Kim W. J., et al., Epigallocatechin Gallate Protects against Hypoxia-Induced Inflammation in Microglia via NF-κB Suppression and Nrf-2/HO-1 Activation, Int J Mol Sci. 7 (23) (2022) 4004.

26. Yang L., Zhang W., Chopra S., et al., The Epigenetic Modification of Epigallocatechin Gallate (EGCG) on Cancer, Curr Drug Targets. 11 (21) (2020) 1099–104.

27. Chen J., Li G., Sun C., et al., Chemistry, pharmacokinetics, pharmacological activities, and toxicity of Quercitrin, Phytother Res. 4 (36) (2022) 1545–75.

28. Lamb Y. N., Rosuvastatin/Ezetimibe: A Review in Hypercholesterolemia, Am J Cardiovasc Drugs. 4 (20) (2020) 381–92.

29. Boutari C., Karagiannis A., Athyros V. G., Rosuvastatin and ezetimibe for the treatment of dyslipidemia and hypercholesterolemia, Expert Rev Cardiovasc Ther. 7 (19) (2021) 575–80.

30. Strilchuk L., Tocci G., Fogacci F., et al., An overview of rosuvastatin/ezetimibe association for the treatment of hypercholesterolemia and mixed dyslipidemia, Expert Opin Pharmacother. 5 (21) (2020) 531–39.

31. Cortese F., Gesualdo M., Cortese A., et al., Rosuvastatin: Beyond the cholesterol-lowering effect, Pharmacol Res. (107) (2016) 1–18.

32. Das Y. T., Bagchi M., Bagchi D., et al., Safety of 5-hydroxy-L-tryptophan, Toxicol Lett. 1 (150) (2004) 111–22.

33. Chen S., Zhao H., Yan X., et al., 5-Hydroxy-l-tryptophan Promotes the Milk Calcium Level via the miR-99a-3p/ATP2B1 Axis in Goat Mammary Epithelial Cells, J Agric Food Chem. 10 (68) (2020) 3277–85.

34. Stumpf J. L., Deferasirox, Am J Health Syst Pharm. 6 (64) (2007) 606–16.

35. Galanello R., Campus S., Origa R., Deferasirox: pharmacokinetics and clinical experience, Expert Opin Drug Metab Toxicol. 1 (8) (2012) 123–34.

36. Yui J. C., Geara A., Sayani F., Deferasirox-associated Fanconi syndrome in adult patients with transfusional iron overload, Vox Sang. 7 (116) (2021) 793–97.

