## Supplemental Figure S1,Figure S2,FigureS3 for "Crystal structure of mRNA cap (guanine-N7) methyltransferase E12 subunit from monkeypox virus and discovery of its inhibitors"

**Supplementary Figures**

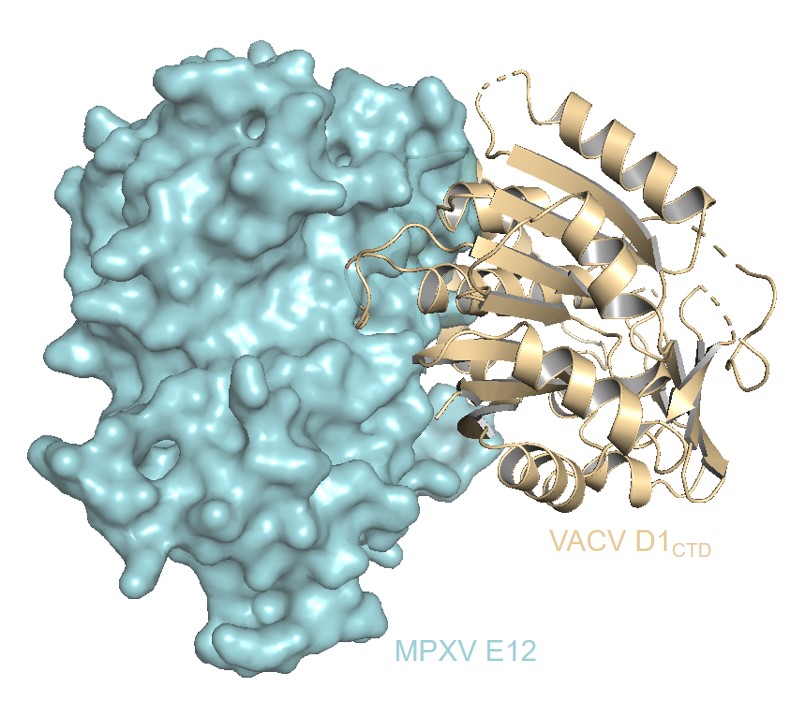

**Fig. S1. The structure of** **E12-D1 complex.** The surface of E12 subunit is shown in green and the D1_CTD_ subunit is colored in gold.

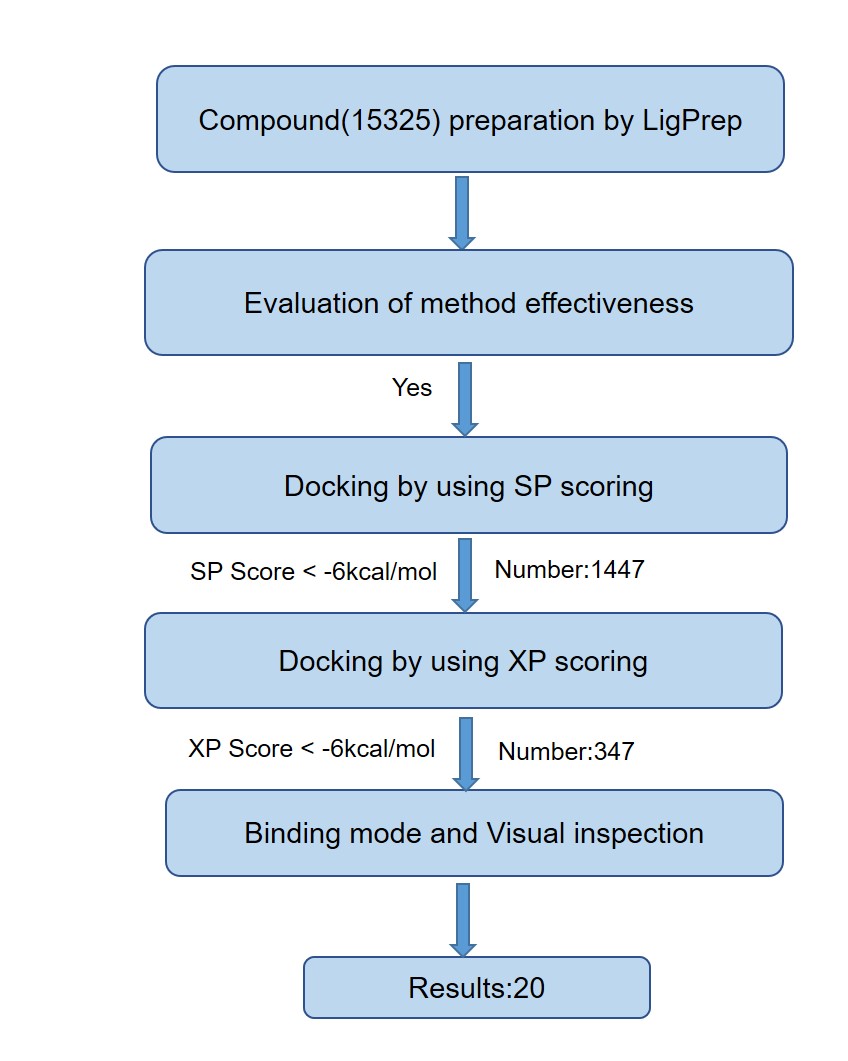

**Fig. S2. The process of virtual screening for molecular docking.** All 15, 325 compounds were prepared according to the default settings of the LigPre module. The ligand was docked to evaluate the effectiveness of the docking method. The dataset was then screened by SP docking. A total of 1,447 compounds had SP score < −6 kcal/moL. The SP docking template is suitable for mass screening of compounds. Next, the XP docking template was used to screen out the pro-ligands with higher scores determined by the SP method and a total of 347 compounds have an XP score < −6 kcal/mol. Finally, 20 compounds were selected by binding mode and visual inspection.

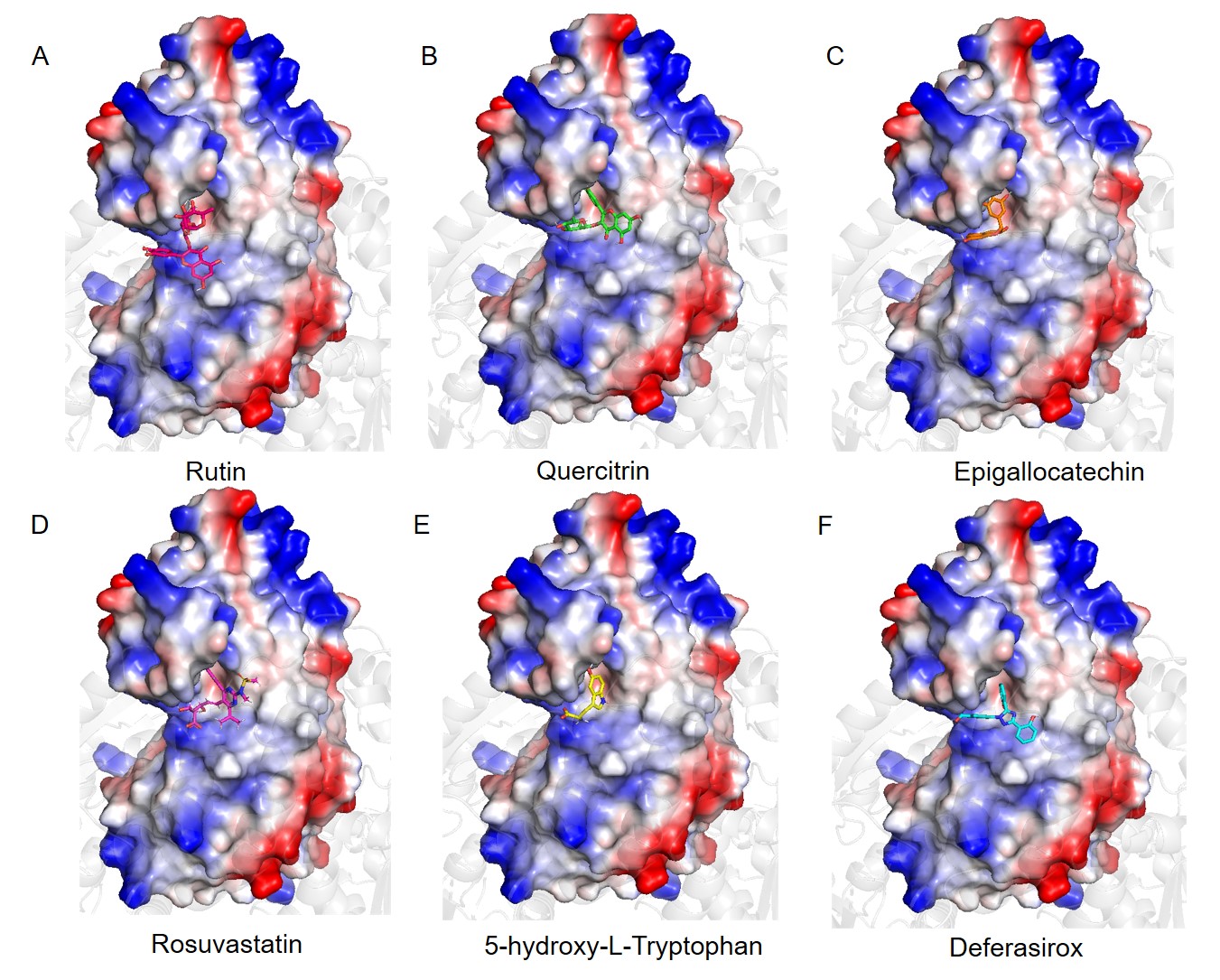

**Fig. S3. The surface charge distribution and the drug-binding sites on the E12 subunit.** The structures of E12 subunit in complex with (A) Rutin, (B) Quercitrin, (C) Epigallocatechin, (D) Rosuvastatin, (E) 5-hydroxy-L-Tryptophan, and (F) Deferasirox. The surface of E12 subunit is shown in vacuum electrostatics.
